# *C. mastitidis* requires the protein Sortase F to colonize the eye

**DOI:** 10.64898/2026.03.12.711320

**Authors:** Yannis E. Rigas, Jackie Shane, Benjamin Treat, Robert M.Q. Shanks, Anthony J. St. Leger

## Abstract

The ocular surface is a mucosal tissue that is constantly exposed to environmental antigens and potential pathogens. Human microbiomes play a critical role in the balance of surveillance and inflammation at sites of colonization. Historically, the investigation of the ocular microbiome has been difficult due to its paucibacterial nature and the inhospitable environment of the ocular surface. Despite this, *Corynebacterium mastitidis* (*C. mast*) developed a unique ability to colonize the eye and elicit a protective immune response characterized by induction of IL-17 from γδ T cells and protection from corneal infection. Therefore, we sought to understand the unique bacterial machinery that *C. mast* utilizes to colonize the eye and how it affects the induction of an eye-specific immune signature. Using a *C. mast* transposon mutant library, we identified a mutant that completely lacked an ability to form biofilm, colonize the eye, and induce *in vivo* immunity. Whole genome sequencing revealed a disruption in the sortase F gene, which anchors proteins to the cell wall of *C. mast*, governing biofilm formation and tethering of adhesins to the cell surface. Additionally, we show that mutation in individual *C. mast* adhesins does not affect ocular colonization or immune induction. By understanding the molecular mechanism of ocular microbial colonization, this work advances our understanding of how bacteria colonize and induce immune responses on the eye, providing a foundation for developing novel therapeutic strategies against ocular infections.

## Introduction

The ocular surface is continuously exposed to environmental antigens including allergens, pathogens, and microbes that have the potential to trigger inflammatory responses detrimental to vision. To protect against infection while maintaining a transparent surface for visual function, the eye has evolved several mechanisms that balance surveillance with inflammation control^1^. These mechanisms include continual washing of tears across the eye, high tear concentrations of anti-microbials, limited nutritional resources, and others^2^. Despite this inhospitable environment, a resident microbial community has been described that contains the genera *Corynebacterium, Staphylococcus, Cutibacterium*, and S*treptococcus*^3^. Building off these studies, *Corynebacterium mastitidis* (*C. mast*) has been identified as an ocular commensal that induces a protective immune signature characterized by IL-17 production from γδ T cells and protection from blinding pathogens^4^. While recent advances have identified *C. mast* factors that stimulate its immune signature, identification of a specific microbial factor governing colonization of the eye is currently undescribed^5^.

Core members of microbiomes have evolved ways to either avoid detection or prevent elimination by host immune defenses. In the gut, components of the microbiome developed ways to thrive in oxygen deficient conditions, break down complex carbohydrates from ingested foods, form biofilms to protect from host defenses, and produce factors that regulate or dampen immune responses and surveillance^6,7^. Similarly, given the harsh environment of the ocular surface, eye-colonizing microbes needed to develop ways to thrive at the ocular surface. Previous studies support the notion that eye-colonizing microbes use biofilm formation to protect themselves from host defenses allowing them to remain at the ocular surface indefinitely^4,8^; however, no study has attempted to assess the impact of biofilm formation on *C. mast*’*s* ability to colonize the ocular surface.

Biofilm formation is often mediated by sortases since they are known to contribute to surface anchored proteins associated with bacterial colonization of other surfaces/tissues^9,10^. Sortases are a class of membrane bound enzymes that cleave secreted proteins through recognition of a binding motif, most commonly LAxTG or LPxTG, and anchor these proteins to the bacterial surface or onto repeated membrane bound chains^11,12^. Surface display of these proteins orchestrated by sortase cleavage contributes to bacterial phenotypes such as attachment to surfaces, virulence, biofilm, and host-microbe interactions^13-16^. Sortases are classified into six classes (A-F), with each having various roles in anchoring surface proteins (A, E), iron acquisition (B), pili polymer formation (C), and display of proteins enabling spore formation (D)^17-20^. While sortases A-E are well characterized, sortase F is less understood. However, sortase F has recently been described in *Propionibacterium acnes* where it acts as sole transpeptidase similar to sortase A^21^; however, explicit functional characteristics of sortase F have not been described. While Corynebacteria harbor sortases^22,23^, we found that *C. mast* possesses a mechanistically and genetically distinct sortase F, suggesting that this sortase may be playing an active role in *C. mast*’s ability to colonize the eye.

Here, we describe a unique sortase F in *C. mast* that governs its ability to colonize the eye long-term allowing for induction of a protective immune signature. This highlights the role for sortases in host-microbe interactions at the ocular surface, which may elucidate how commensals, and pathogenic bacteria, may colonize this inhospitable environment. This research could be utilized for the development of novel therapeutics to treat ocular infections.

## Results

### C. mast sortase F is necessary for biofilm formation and ocular colonization

The ocular surface maintains a paucibacterial microbiome due to harsh environmental conditions that restrict microbial colonization^24,25^. Notably, *C. mast* is able to overcome these barriers and colonize the ocular surface while inducing an immune signature that protects the eye from pathogenic infection^4^. While this relationship is known, the microbial factor(s) that govern *C. mast*’s ability to colonize the eye remains elusive. Thus, we created a transposon mutant library that generated thousands of single insertion mutants that could be used to screen for factors related to colonization and immune induction(**Figure S1A**)^26^. Because biofilm formation is known to contribute to attachment of microbes on surfaces, we hypothesized that *C. mast’*s ability to form a stable biofilm contributes to its colonization and immune induction at the ocular surface^27,28^. Thus, we performed a high throughput screen of our whole *C. mast* transposon mutant library and identified a mutant (*SortF::Tn*) that does not produce a stable biofilm when tested *in vitro* (**Figure 1A& S1B**).

**Figure 1:**
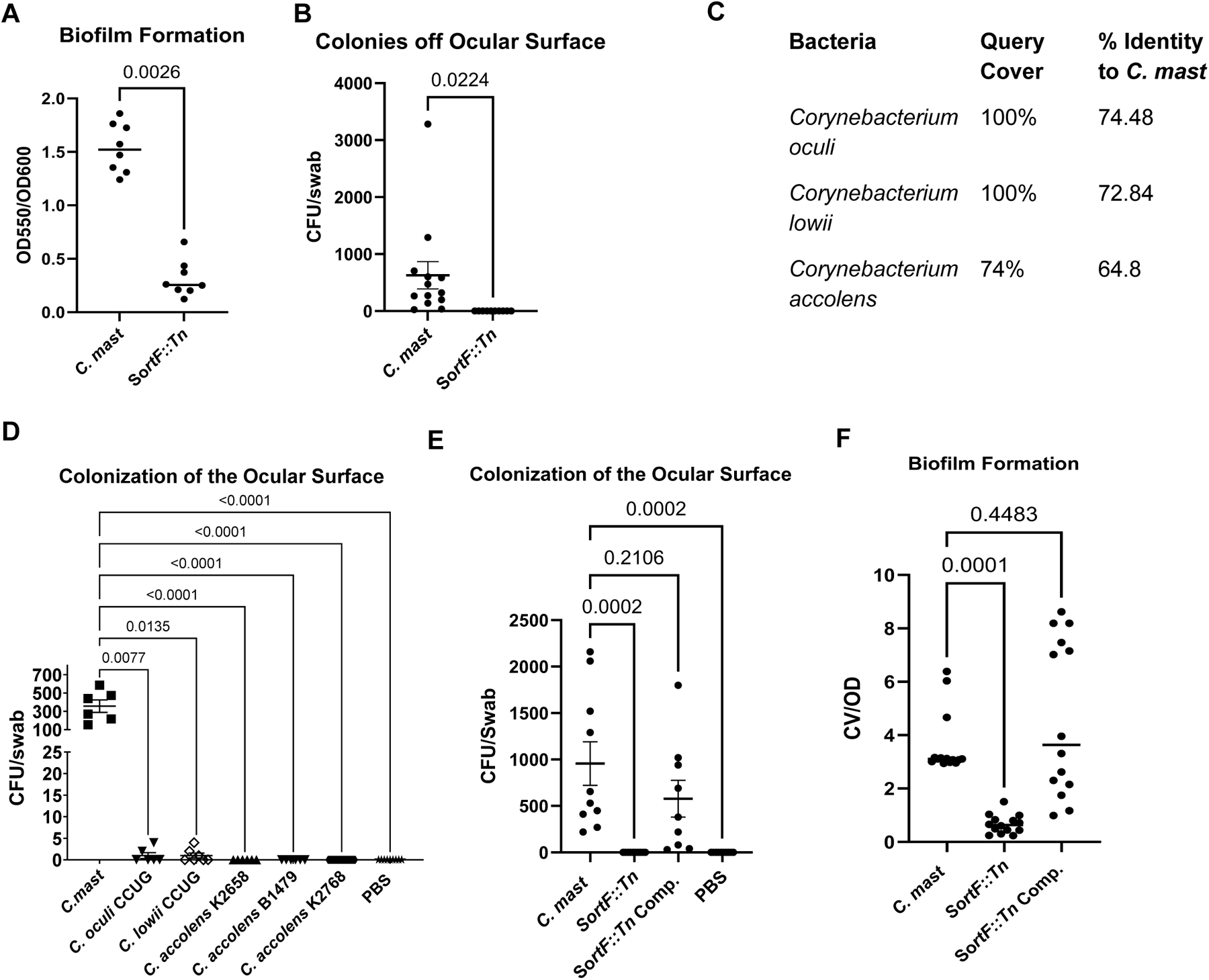
*C. mast* sortase F is necessary for biofilm formation and ocular colonization. (A) *C. mast* or mutant *SortF::Tn* were grown in 48-well tissue culture plates for 48 hours. Bacterial growth was quantified at OD_600_ (OD), wells were washed, and biofilm was stained with 0.1% crystal violet and quantified at OD_550_. Graph shows the amount of biofilm normalized to growth (OD_550_/OD_600_), where symbols represent individual experiments. Significance was determined using a Kruskal-Wallis test. (B) Three-week-old C57BL/6 mice were inoculated every other day for a total of three inoculations with 5×10^5^ CFU/eye of WT *C. mast* or *C. mast* mutant *SortF*::Tn. One week post final inoculation, eyes were swabbed and streaked on LBT plates with Fosfomycin. Graphs show the number of CFU/swab, where symbols represent individual mice from three independent experiments. Significance was determined using a Welch’s T test. (C) Bacterial DNA was isolated and sent to the Microbial Genome Sequencing Center for whole-genome sequencing. and sequences were compared to other corynebacterial using BLASTP. (D) As described in B, mice were inoculated with WT *C. mast*, clinical isolates of *C. accolens*, or commercially purchased *C. oculi and C*.*lowii*. A week after the final inoculation, eyes were swabbed and bacterial colonization was quantified. Graphs show the number of CFU/swab, where symbols represent individual mice from three independent experiments. Significance was determined using a Welch’s T test. (E) As described in B and E, mice were inoculated with WT *C. mast, SortF::Tn*, or complemented *SortF::Tn*. A week after the final inoculation, eyes were swabbed and bacterial colonization was quantified. Graphs show the number of CFU/swab, where symbols represent individual mice from three independent experiments. Significance was determined using an ordinary one-way ANOVA. (F) As described in A, *C. mast, SortF::Tn*, or *SortF::Tn* Complement were grown in 48-well tissue culture plates for 48 hours. Bacterial growth was quantified at OD_600_, wells were washed, and biofilm was stained with 0.1% crystal violet. Graph shows the amount of biofilm normalized to growth (OD_550_/OD_600_), where symbols represent individual experiments. Significance was determined using an ordinary one-way ANOVA.

To better understand the physiological consequences of transposon insertion in our mutant, we whole genome sequenced our mutant and identified that its transposon insertion was within its *Sortase F* gene. In order to describe how sortase F contributes to *C. mast*’s unique ability to colonize and induce protective immunity, we attempted to colonize the ocular surface of mice and found that *SortF::Tn* is unable to colonize the eye (**Figure 1B**), even though it grows similarly to WT *C. mast* in culture. We then compared the peptide sequence of *C. mast* Sortase F to other *Corynebacteria*, revealing that *C. mast*’s sortase F is unique among *Corynebacteria*, potentially explaining *C. mast*’s distinct ability to efficiently colonize the mouse eye (**Figure 1C**). Further, inoculation with *Corynebacteria* containing the most similar sortase F protein sequences showed limited to no colonization ability (**Figure 1D**), suggesting that the unique features of *C. mast*’s sortase F drives its ability to colonize and persist on the ocular surface. To confirm that sortase F drives this biofilm formation and subsequent colonization of the eye, we generated a complemented strain of *SortF::Tn* with WT sortase F expression. Using this complemented mutant, termed *SortF::Tn Comp*., we demonstrated a rescue of both biofilm formation (**Figure 1E**) and colonization phenotypes (**Figure 1F**). Confirming that sortase F in *C. mast* is unique and necessary for both biofilm formation and ocular surface colonization.

### C. mast sortase F tethers adhesins to cell surface

The molecular mechanisms of sortase F function in *C. mast* are currently undescribed. Having demonstrated that functional sortase F is necessary for biofilm formation and ocular colonization in *C. mast*, we next investigated its specific molecular role. Sortases are known in other bacteria and *Corynebacteria* to tether membrane bound proteins to the bacterial cell surface, suggesting a potential mechanism for *C. mast* sortase F function. To test this, we first sorted the *C. mast* proteome for secreted and cell wall proteins using SignalP5^29^. Additionally, all non-cytoplasmic proteins were analyzed for the standard LAxTG and LPxTG sortase binding motif using the FIMO^30^ software of MEME suite^31^. However, no proteins were found that contained these motifs, thus, we further analyzed the membrane proteomes of WT *C. mast* and mutant *SortF::Tn* for differences using mass spectrometry. These analyses revealed a nearly complete loss of the sortase F protein along with significantly decreased adhesin 1 (Adh1) and 2 (Adh2) surface proteins (**Figure 2A**). Indicating that expression of sortase F in *C. mast* is necessary for proper cellular trafficking of adhesins on *C. mast*’s surface. To validate that these adhesins were indeed sortase substrates we analyzed their protein sequence using GPApred^32^. Both adhesin 1 and 2 came back as 100% predicted sortase substrates despite their lack of standard motif. We next asked whether the lack of adhesins in *SortF::Tn* or sortase F itself is responsible for the loss of biofilm formation and colonization using *Adh1::Tn* and *Adh2::Tn* mutants. These studies showed that Tn disruption of Adh1 impairs *C. mast*’s ability to produce a biofilm, while Adh2 disruption does not (**Figure 2B**). Additionally, neither Adh1 nor Adh2 is necessary for *C. mast* colonization (**Figure 2C**), revealing that sortase F itself, not individual downstream protein targets, is necessary for *C. mast*’s unique ability to colonize the eye. To demonstrate that sortase F is responsible for decreased adhesin expression on *C. mast*’s surface, we performed trypsin shaving of cell surface proteins on our mutants and compared them WT and the complimented strains. *Adh2::Tn* was excluded from analysis due to poor cell pellet formation. This analysis revealed that Tn disruption of Adh1 does not influence protein abundance on the *C. mast* surface, while complementation of *SortF:Tn* rescues the loss of cell surface proteins (**Figure 2D**). Altogether, these data illustrate that sortase F in *C. mast* is necessary for adhesin expression on the cell surface, and while these adhesin proteins contribute to biofilm formation, there is likely redundancy between the two adhesins in *C. mast’s* ability to colonize the ocular surface.

**Figure 2:**
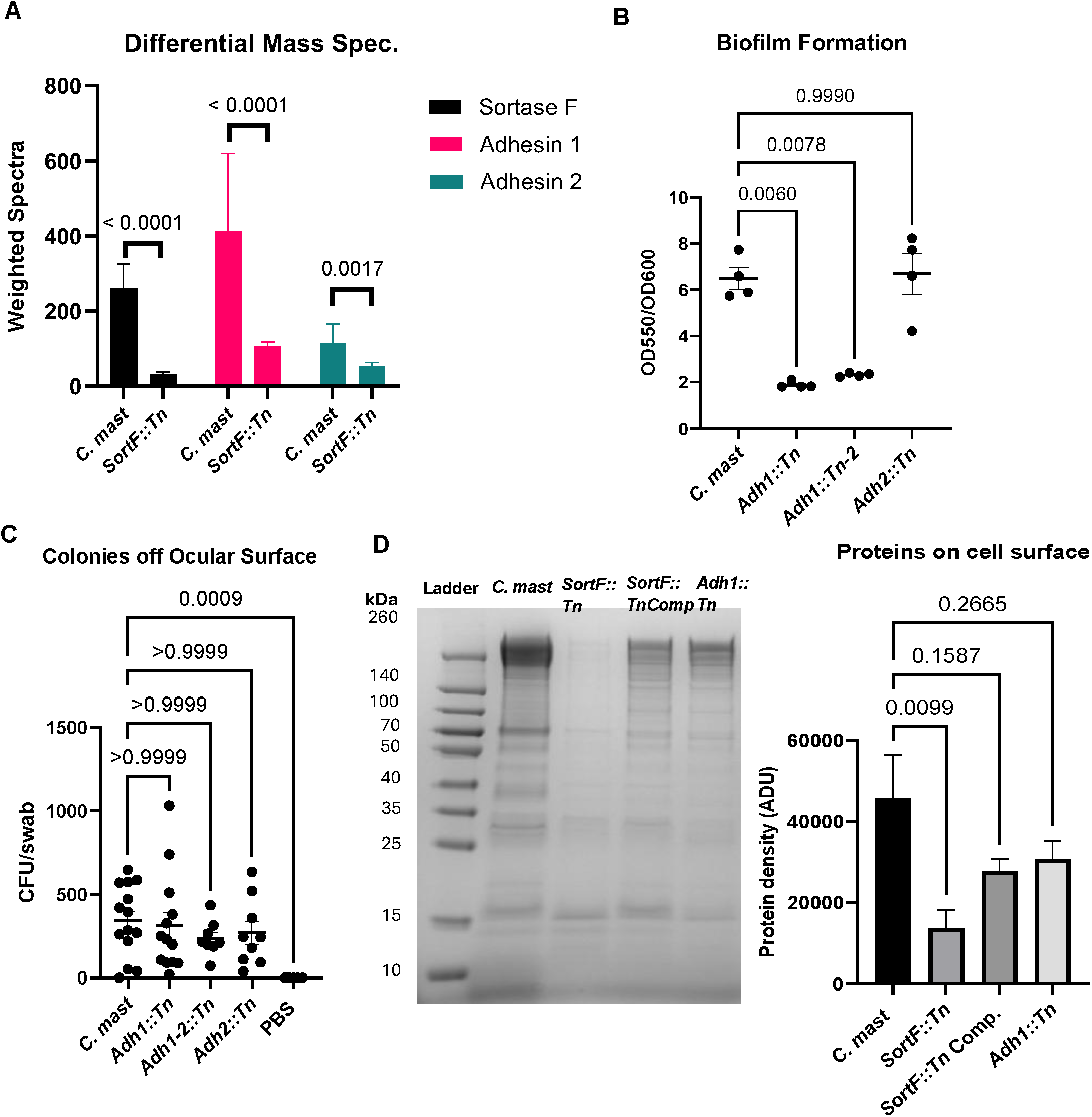
*C. mast* tethers adhesins to cell surface. (A) WT *C. mast* and *SortF::Tn* cell wall associated proteins were shaved off using trypsin digestion and sent for mass spectrometry (MS Bioworks). Weighted spectra abundance from three independent replicates for each sample were evaluated using Fisher’s Exact Test (B) As described in Figure 1A, WT *C. mast, Adh1::Tn, Aap::Tn, Adh1-2::Tn*, and *Adh2::Tn* were grown in 48-well tissue culture plates for 48 hours, bacterial growth was quantified at OD_600_, wells were washed, and biofilm on was stained with 0.1% crystal violet and quantified at OD_550_ after resuspension in ethanol. Graph shows the amount of biofilm normalized to growth (OD_550_/OD_600_), where symbols represent individual experiments. Significance was determined using a Brown-Forsythe and Welch ANOVA test. (C) Three-week-old C57BL/6 mice were inoculated every other day for a total of three inoculations with 5×10^5^ CFU/eye of WT *C. mast, SortF*::*Tn, Adh1::Tn, Aap::Tn*, or *Adh1-2::Tn*. One week post final inoculation, eyes were swabbed and streaked on LBT plates with Fosfomycin. Graphs show the number of CFU/swab, where symbols represent individual mice from three independent experiments. Significance was determined using a Kruskal-Wallis test. (D) As described in A, bacterial cell walls were shaved using trypsin and products were run on an SDS page gel. Protein density per lane was quantified using FIJI. Significance was assessed by standard one-way ANOVA

### C. mast adhesins behave similarly to previously described adhesins in other species

To better understand the characteristics of these *C. mast* adhesins, we predicted their structures using AlphaFold3^33^. This analysis revealed that these adhesins follow the standard IgG-like fold arrangement typical of bacterial adhesins, with additional repeats generating larger molecular structures **(Figure 3A&B)**. These adhesins are composed of highly structured β-barrel repeats with terminal α-helices connected by unstructured linkages. Since adhesins with similar structures are known to bind extracellular matrix components such as collagen and fibronectin^34^, we tested whether these mutants could bind these molecules *in vitro*. Binding assays revealed that *SortF::Tn, Adh1::Tn*, and *Adh2::Tn* were all significantly reduced in their ability to bind collagen (**Figure 3C**), while only SortF::Tn showed significantly decreased fibronectin binding (**Figure 3D**). These results suggest that disruption of sortase F in *C. mast* impairs its ability to bind ocular surface molecules, contributing to its loss of colonization.

**Figure 3:**
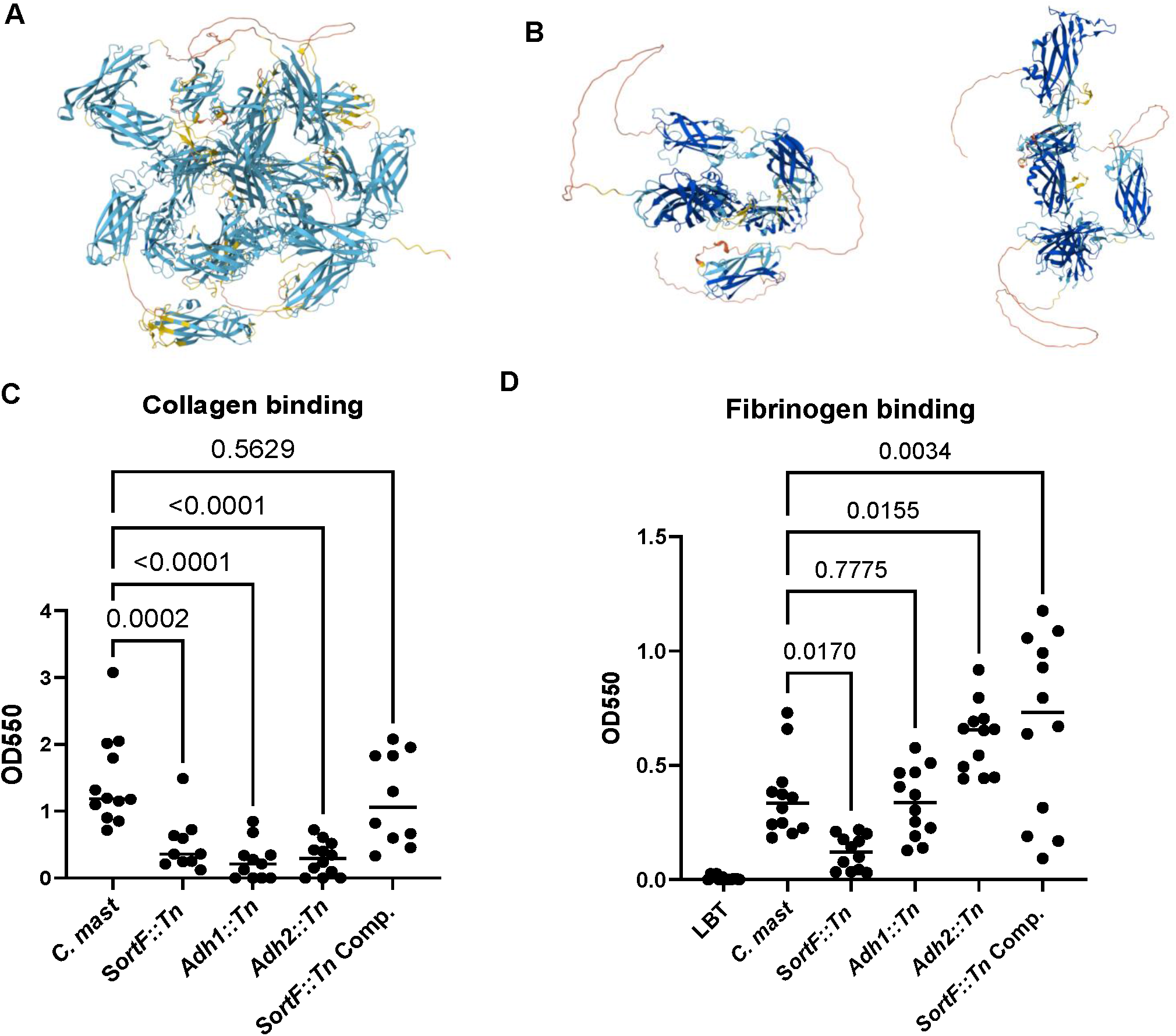
*C. mast* adhesins behave similarly to previously described adhesins in other species. (A&B) Peptide sequences of both adhesins were analyzed using Alphafold3. Both show the standard IgG-C Beta barrel structure connected by unstructured linkages (A) Adhesin 1 shows a more globular structure while (B) Adhesin 2 is more flattened with obviously less repeats of the IgG-C beta barrels.(C&D) The ability for the transposon mutants to bind collagen and fibrinogen after 4 hours of growth on ELISA plates coated with (C) 100μg/mL collagen type1 or (D) 10 μg/mL of fibrinogen.

### Colonization is necessary for protective immune signature on the eye

Colonization with *C. mast* induces a protective immune signature characterized by recruitment of γδ T cells to the conjunctiva and downstream signaling that recruits neutrophils and enhances anti-microbial production while causing no pathology in immunocompetent hosts^4^. Thus, we asked whether *SortF::Tn* is able to induce a similar immune signature even though it does not colonize. Using our *in vivo* model, we show that *SortF::Tn* does not recruit IL-17 producing γδ T cells to conjunctiva, and, downstream of IL-17, recruit neutrophils (**Figure 4A&B**). While these results show that *SortF::Tn* cannot induce an ocular immune signature similar to WT *C. mast*, this does not definitively show that it lacks the immunostimulatory components necessary for IL-17 induction from γδ T cells. Using an *in vitro* stimulation model, we tested the ability of *SortF::Tn* to stimulate IL-17 production from γδ T cells and showed that it is able to stimulate IL-17 similarly to WT *C. mast*. These findings demonstrate that while sortase F is required for *C. mast* colonization, it does not regulate expression of *C. mast*’s immunostimulatory components (**Figure 4D**) like trehalose monocorynomycolate or conventional TLR2 ligands such as lipoteichoic acid^5,35^. To fully describe the role of sortase F in *C. mast* ocular establishment and immune phenotype, we asked whether sortase F is essential for initial establishment of *C. mast* colonization and/or is it necessary for long term stability. We have previously shown that inoculation with a component of the *C. mast* lipid membrane is able to induce its typical immune signature^36^ suggesting that daily topical application of γδ T cell stimulants is sufficient to induce tissue-specific immunity. Using this same approach, we topically applied *SortF::Tn* to the eye daily for seven days. After 7 days, we assessed the conjunctival γδ T cell response. Notably, we observed that *SortF::Tn* failed to recruit Vγ4 γδ T cells to the eye (**Figure 4E**), signifying that sortase F is necessary for initial establishment of colonization and subsequent induction of *in vivo* immunity. Overall, we have shown that sortase F is essential for *C. mast* to produce biofilm and express adhesin proteins on its surface which contributes to *C. mast*’s ability to establish colonization on the ocular surface and subsequent immune signature.

**Figure 4:**
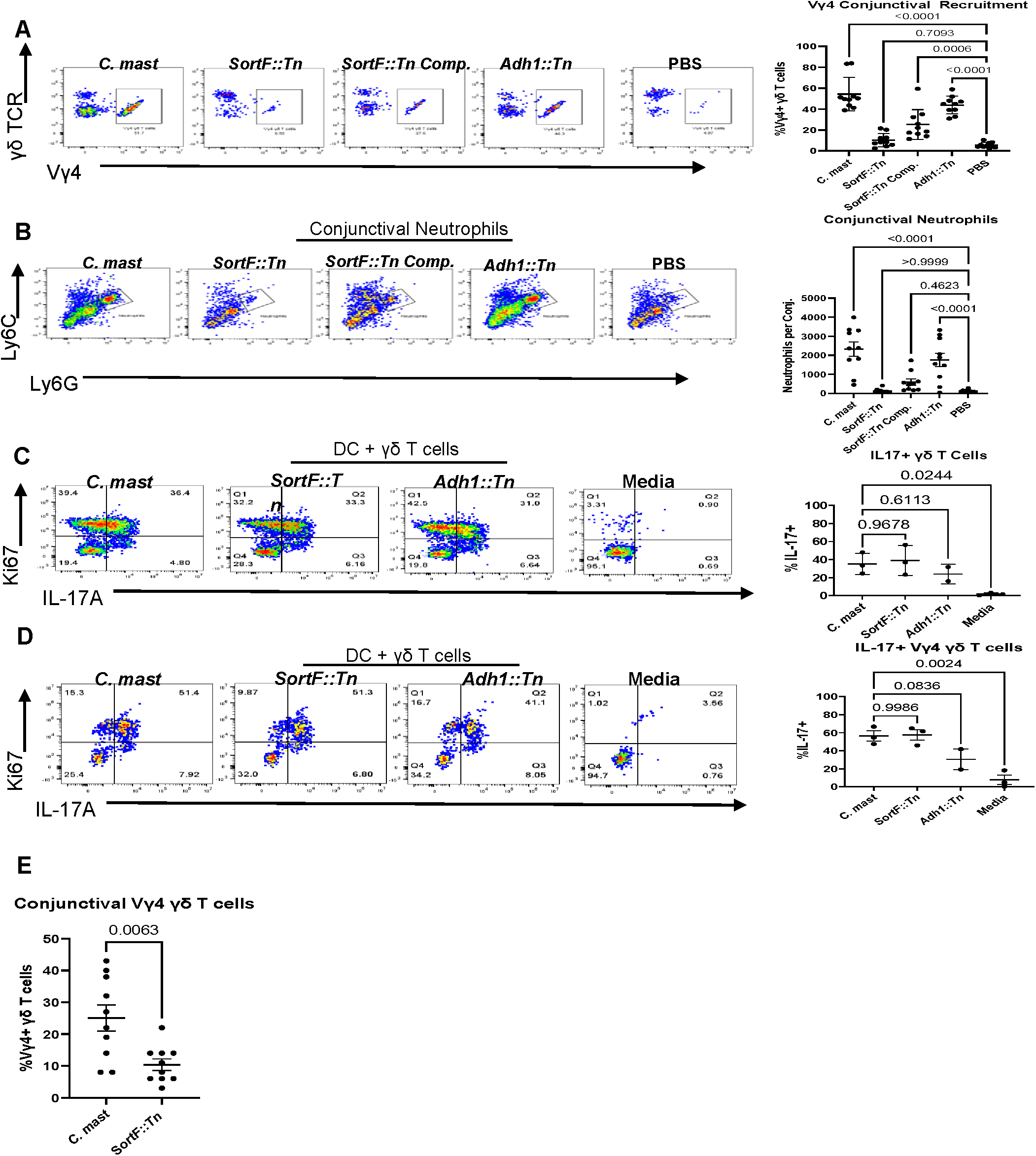
Colonization is required for induction of protective immunity on the ocular surface. (A&B) Two weeks after the final inoculation of *C*.*mast* and mutants, mice were sacrificed, and conjunctivas were harvested. Single cell suspensions were stimulated with PMA/Ionomycin and analyzed for (A) IL-17+ Vγ4 γδ T cells; (B)conjunctivas were also assessed for number of neutrophils by flow cytometry. (A&B) Symbols represent data for individual mice from three independent experiments. Bars represent average ± SEM. Significance was determined using and ordinary one-way ANOVA. (C&D) 1×10^4^ γδ T cells were obtained from WT mice which pre-associated with *C. mast* and cocultured with 1×10^5^ DCs pretreated with 1×10^5^ CFU of either WT *C. mast, SortF::Tn*, or *Adh1::Tn* 24 hours. After 72 hours of coculture, γδ T cells were assessed for percent of IL-17+Ki67+ γδ T cells and Vγ4 γδ T cells by flow cytometry. Bars are mean ± SEM. Symbols represent individual experiments. Significance was determined using an ordinary one-way ANOVA. (E) Three-week-old C57BL/6 mice were inoculated every day for 7 days with 5×10^5^ CFU/eye of WT *C. mast* or *SortF*::*Tn*. The day after the final inoculation, conjunctivas were harvested and recruitment of γδ T cells was assessed using flow cytometry. Symbols represent data for individual mice from two independent experiments. Bars represent average ± SEM. Differences were determined to be significant using a Welch’s t test.

## Discussion

The contribution of ocular resident microbes to ocular health and disease has recently become a research topic of interest, with studies showing how ocular microbes induce a protective immune signature that inhibits pathogenic infection^4^. While recent studies have uncovered immunostimulatory molecules driving induction of protective immunity, a conserved microbial factor that regulates colonization of the ocular surface has remained elusive^5^. In this study, we identify a novel sortase F in *C. mast* that contributes to biofilm formation by tethering adhesins to the cell wall enabling *C. mast* to uniquely colonize the ocular surface. Despite the requirement for sortase F in the colonization of the ocular surface, functional sortase F is not required for the *in vitro* induction of IL-17. Therefore, we can conclude that sortase F does not regulate the production of γδ T cell stimulants, which we have previously shown to be lipoteichoic acid and TMCM^5,35^. Alternatively, we conclude from our collective data that functional sortase F is critical for colonization of the ocular surface. Notably, even repeated exposure of the sortase F deficient *C. mast* mutant does not induce protective immunity. Therefore, we concluded that sustained colonization is a requisite for induction of lasting immunity at the ocular surface. Collectively, our work establishes sortase F as a critical determinant of ocular commensal colonization by *C. mast* and therefore the establishment of protective immunity. Further, our data provide a foundation for understanding how microbial factors govern tissue-specific colonization and may open avenues for developing sortase-based therapeutics to modulate ocular immunity and control of infections.

## Materials and Methods

### Bacterial culture and genetic constructs

WT *C. mast* and all mutants were grown in Luria-Bertani (LB) liquid medium supplemented with 0.5% Tween-80 (LBT). Cultures were supplemented with 100 μg/mL fosfomycin for Corynebacterium selection, and mutant cultures received an additional 50 μg/mL kanamycin for transposon selection. Bacteria were streaked on 1.5% agar LBT plates. All bacterial cultures were grown at 37°C with liquid cultures maintained under constant agitation.

### Creation of *C. mast* mutants and Sortase F complementation plasmid

As previously described, *SortF::Tn, Adh1::Tn, Adh2::Tn* were generated in previously generated transposon mutant libraries^26,37^. Additionally, we generated a *C. mast* sortase F complementation plasmid from a previously described plasmid, and transforming *SortF::Tn* through electroporation and antibiotic selection^37^.

### Mice

Female C57BL/6 mice purchased from Jackson Laboratories (Bar Harbor, ME, USA) were housed in the Animal Resource Facility at the University of Pittsburgh Medical Center (Pittsburgh, PA, USA) and colonized with *C. mast* at 3 weeks of age in all experiments according to our previous inoculation schemes^4,5^. The use of animals was in accordance with the ARVO Statement for the Use of Animals in Ophthalmic and Vision Research. University of Pittsburgh IACUC approval number: 23063056

### Characterization of *SortF::Tn*

We identified *SortF::Tn*, a mutant unable to produce biofilms *in vitro*, by performing a high throughput screen for biofilm formation using a crystal violet based assay^38^. Further, *SortF::Tn* and WT *C. mast* were grown in a 96 or 48 well tissue culture treated dish at 37°C for 48 hours. After incubation, growth was quantified at OD_600_, supernatants were removed, and wells were washed three times with DI water. Biofilm was then stained with 100 μl of 0.1% crystal violet and incubated at room temperature for 15 minutes. After incubation, wells were washed three times with DI water, and 100 μl of 190 proof ethanol was added to the well and incubated at room temperature for 15 minutes. Plate was then analyzed by quantifying absorbance in a plate reader at 550 nm and the amount of biofilm was normalized to growth.

C57BL/6 mice were inoculated every other day for a total of three inoculations with 5×10^5^ CFU/eye of WT *C. mast* or *SortF::Tn*. One week post-final inoculation, eyes were swabbed and streaked on LBT plates with Fosfomycin.

### Discovery of Sortase substrates

To elucidate which proteins are substrates for sortase in *C. mast*, we shaved off proteins from the cell surface then performed differential mass spectrometry. First, we grew WT *C. mast* and *SortF::Tn* to log phase overnight and diluted both cultures to 0.5 OD_600_. One milliliter of each culture and an LBT control were subjected to trypsin shaving as described in Bonn et all, 2018^39^. These samples were run on Bolt 12%, Bis-Tris Plus Mini Protein gels from Invitrogen to confirm protein presence in sample and to qualitatively assess differences between WT *C. mast* and *SortF::Tn*. These samples were then sent for differential mass spectrometry through MS Bioworks. The spectra output was assessed using Scaffold 5 and the weighted spectra were analyzed for significance using Fisher’s exact test with benjamini-hochberg correction.

### Characterization of Sortase substrates

After finding that both substrates predicted in the differential mass spectrometry had been interrupted in previous transposon libraries, we used these mutants to try to understand the function of each protein better. First, we performed a biofilm assay with both mutants and colonized mice with both mutants as described previously. We also extracted cell wall-associated proteins as described previously and ran them on the mini protein gels; unfortunately, the cell debris never pelleted properly from the Adhesin 2 KO, PhoZ 21 so that was excluded. The density of protein in each lane was measured using FIJI and analyzed in Prism.

Next the function of these proteins was analyzed using a collagen and fibrinogen binding assay. For both methods a Corning Costar 96 well high binding assay plate was coated with 100 µl of either 100 µg/mL rat tail Collagen I or 10 µg/mL mouse fibronectin (Fisher scientific) overnight at 4 ºC. The plates were washed 3 times with PBS with 0.05% tween 80 then blocked with 2% BSA for 30 minutes and the wash was repeated. Log phase overnight cultures of bacteria were diluted to 0.5OD_600_ before 100µl was added to the wells of the precoated plates. They were incubated for 4 hours at 37ºC then washed and 100µl of 1% gentian violet was added to each well. After 15 minutes the wells were washed gently three times and let to dry overnight. Once dry the crystal violet was solubilized with 50/50 ethanol and acetic acid before the absorbance was read on a spectrophotometer. Lastly, the protein sequence of each adhesin was run through alphafold3^33^.

### *in vivo* Immune Signature Assessment

Three weeks after the final inoculation of WT *C. mast* or our mutants, mice were sacrificed by cardiac perfusion and conjunctivae were isolated as previously described^4,5,26^. Briefly, tissue was excised, minced, and incubated in a collagenase solution at 37°C for one hour. After incubation, tissue was passed through a Corning Falcon Test Tube with Cell Strainer Snap Cap (352235; Corning Inc., Corning, NY, USA). Tissue was then stained for neutrophil analysis, or stimulated with phorbol myristate acetate (PMA)/ionomycin in the presence of brefeldin A for 4 hours at 37°C. After the 4-hour incubation, samples were stained for Vγ4 TCR and IL-17A. All samples were analyzed using CytoFLEX LX Flow Cytometer and FlowJo v10.8.

### *in vitro* Immune Assessment

1×10^5^ bone marrow derived dendritic cells (BMDCs) were pulsed for 24 hours with 1×10^5^ CFU of *C. mast* or Tn mutants at 37°C and 5% CO_2_. The next day, γδ T cells were isolated from the cervical and inguinal lymph nodes of WT B6 mice using FACS. After isolation, at least 1×10^4^ γδ T cells were incubated with the pre-pulsed BMDCs for 72 hours. Then, cells were harvested and stained with fluorescently labeled antibodies against the γδ TCR and intracellular IL-17A. Samples were analyzed on the CytoFLEX LX Flow Cytometer.

## Notes

### Competing Interest Statement

The authors have declared no competing interest.

### Funding

- NIH: R01EY032482, U24EY035102, T32EY017271, EY036686
- CORE Grant: P30 EY08098

