## Supplemental Figure 1 for "*C. mastitidis* requires the protein Sortase F to colonize the eye"

A

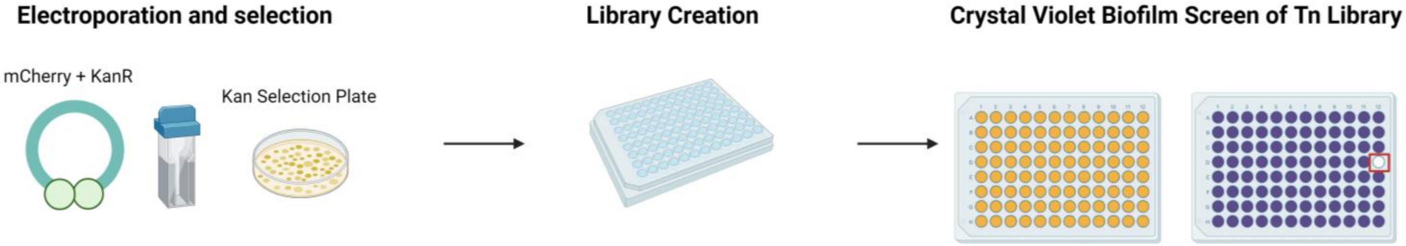


B

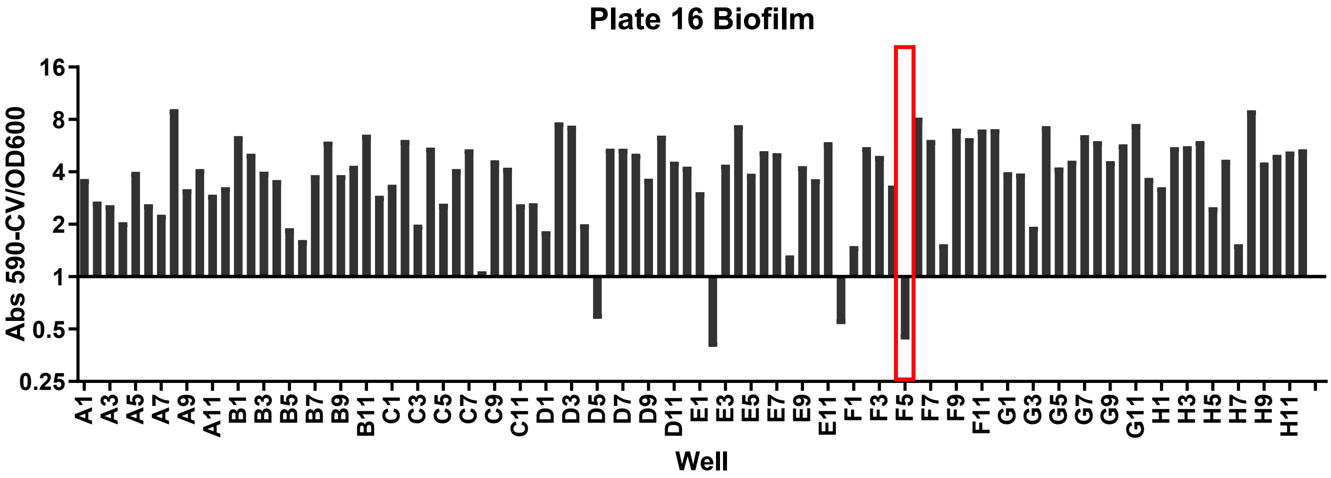


**Figure S1: Identification of *SortF::Tn* using transposon mutant library**

(**A**) Experimental design to create transposon mutant library and perform a high throughput screen for a *C. mast* biofilm knockout. (**B**) A 96 well plate of a *C. mast* transposon mutant library was grown in a tissue culture treated plate for 48 hours. Bacterial growth was quantified at OD600 (OD), wells were washed, and the biofilm on the plate surface was stained with 0.1% crystal violet and quantified at OD550 (CV) after resuspension in ethanol. Graph shows the amount of biofilm normalized to growth (OD550/OD600).
